# *Pseudomonas syringae* evades phagocytosis in animal cells through type III effector-mediated inhibition of the LIM kinase-cofilin system

**DOI:** 10.1101/287508

**Authors:** Sung-Jin Yoon, Soohyun Lee, Jun-Seob Kim, Sang-Hyun Lee, Song Choi, Jeong-Ki Min, Inpyo Choi, Young-Jun Park, Choong-Min Ryu

**Author notes:** Corresponding author. (Y.-J. P.); (C.-M. R.).

## Abstract

Certain animal and plant pathogenic bacteria have developed virulence factors (including effector proteins) that enable them to overcome host immunity. A plant pathogen, *Pseudomonas syringae* pv. tomato (*Pto*), secretes a large repertoire of effectors into plant cells via a type III secretory apparatus, thereby suppressing plant immunity. Here, we show that exposure to *Pto* caused sepsis in mice. Surprisingly, the effector HopQ1 disrupted phagocytosis by inhibiting actin rearrangement via a direct interaction with the LIM domain of the animal target protein LIM kinase, a key regulator of actin polymerization. The results provide new insights into cross-kingdom pathogenicity of bacteria. The current studies demonstrate that certain plant pathogenic bacteria such as *Pto* can be fatal in animals due to cross-kingdom host immune suppression.

## INTRODUCTION

In nature, bacterial infection normally shows a tightly regulated host specificity in which the ability to colonize is dependent on species-specific molecular interactions that determine the host range of the pathogen (e.g., porin proteins, type IV pili, and proteases) (*1*). Despite this barrier, some bacteria show cross-kingdom pathogenicity (*2–4*). To infect hosts across kingdoms, first, a pathogen must evade or overcome the immune systems of different hosts long enough to colonize. The innate immune system (non-antigen-specific immune system) in host-organisms plays an essential role in host defense and is the first hurdle that pathogenic bacteria must overcome (*5*).

Various phagocytes, such as macrophages, neutrophils, and dendritic cells are involved in the mammalian innate immune system. Phagocytosis is triggered upon recognition of pathogens by surface receptors, after which engulfed pathogens are digested within phagolysosomes (*6, 7*). Clearance of infected pathogens is a well-orchestrated process that involves various processes, including recognition, internalization, phagosome formation/digestion, and elimination of debris (*6, 8*).

Phagocytosis, especially internalization, requires (de)polymerization of actin for bacteria to be engulfed. This includes formation and closing of the phagocytic cup by membrane protrusion and retraction (*9*). In this process, (de)phosphorylation of cofilin1 (*10*), a ubiquitous and abundant actin-binding factor that plays an important role in actin dynamics by regulating polymerization/depolymerization (*11*). Innate immune responses and thus, host defenses, depend on the regulation of actin by cofilin1 (*12*). LIM domain kinase 1 (LIMK1) is a serine/threonine protein kinase that also plays an essential role in regulating actin filament dynamics10. Overexpression of LIMK1 promotes the growth, metastasis, and migration of multidrug-resistant osteosarcoma cells (*13, 14*). LIMK1 regulates (de)phosphorylation of cofilin1 therefore, LIMK1 and internalization step of phagocytes are directly linked (*15*). Some animal pathogenic bacteria regulate actin dynamics in phagocytes directly or indirectly, which is one mechanism through which bacteria can evade the mammalian immune system (*16, 17*).

Similar to the importance of cofilin-mediated actin rearrangement in the mammalian innate immune system, the actin (de)polymerization factor-based cytoskeleton network plays a critical role in plant immunity (*18*). In plants, actin-associated proteins facilitate the growth, breakdown, and rearrangement of cytoskeleton, which perform specific immune-related functions (*19, 20*). The growth of plant pathogenic microbes in the intercellular space is suppressed by defense components such as pathogenesis-related (PR) proteins and phytoalexins secreted via the actin-based cytoskeleton network (*21*). Various plant pathogens harbor virulence factors that specifically target the host cytoskeleton including actin dynamics (*19*). Disruption of this network results in impaired secretion of PR proteins and phytoalexins and, consequently, lowers the plant’s defense against infection. Thus, disrupting host immunity by regulating actin dynamics via specific bacterial effectors is a common virulence mechanism used by both plant and human pathogens.

Due to the difficulties involved in studying pathogens in different kingdom hosts, few studies have examined the activity of effectors from plant pathogenic species in animal cells; therefore, there is no evidence that plant pathogenic bacterial effectors also target the actin filaments system in animal cells. Initially, we questioned that if actin-filaments network plays an important role in both animal and plant innate immune system and, if pathogen regulates actin filament dynamics for immune system disruption, it might be possible that plant pathogen regulates the phagocytosis in animal host. Here, we demonstrate that inoculating mice with the plant pathogenic bacteria *Pseudomonas syringae* pv. tomato (*Pto*), which is well-defined plant pathogen to study interactions between host and bacteria (*22*), caused fatal sepsis. Also, we found that the type III effector HopQ1, which is secreted by *Pto,* interacted with the mammalian target protein LIMK1 and inhibited phagocytosis, thereby suppressing the mammalian immune response. These results indicate that HopQ1 controls actin filament dynamics by regulating cofilin1 activity via phosphorylation of LIMK1. These data provide the first evidence of a molecular mechanism by which plant pathogenic bacteria disrupt immune responses in animal cells.

## RESULTS

### *Pseudomonas syringae* suppresses an early step in macrophage-mediated phagocytosis

Because of the striking similarities between the innate immune systems of animals and plants (*23–25*), we wanted to determine whether a plant pathogen could attenuate the immune response of an animal cell. We used flow cytometry and confocal microscopy to examine infection of murine peritoneal macrophages by two green fluorescent protein (GFP)-tagged plant pathogens [*P. syringae* pv. tomato (*Pto*-GFP) and *P. syringae pv.* tabaci (*Pta*-GFP)]. Interestingly, in 2-hour incubation the number of GFP-positive macrophages in the presence of *Pto*-GFP was significantly lower than that in the presence of the other plant pathogen (*Pta*-GFP), or in response to *E. coli*-GFP employed as a control (Fig. 1*A*). Confocal microscopy images also clearly indicated that *Pto*-GFP (green) was less able to be engulfed by macrophages (blue) than the other bacterial strains (Fig. 1*B*). Since flow cytometry signals and plain images from confocal microscopy did not provide information on the subcellular localization of plant pathogens, Z-stack images (X-Z plane) of the *Pta* or *Pto*/macrophage complexes were reconstructed. The images showed that, unlike *Pta*-GFP, *Pto*-GFP attached to the surface of the macrophages rather than internalizing (Fig. 1*C*, white arrows). Additionally, a gentamycin protection assay (*26*) revealed that the macrophages internalized the *Pto* strain to a lesser extent than *Pta* or *E. coli*. The number of internal bacterial cells was determined as the number of colony forming units (CFU) after gentamycin treatment to remove surface-attached bacteria (Fig. 1*D*).

**Fig. 1.**
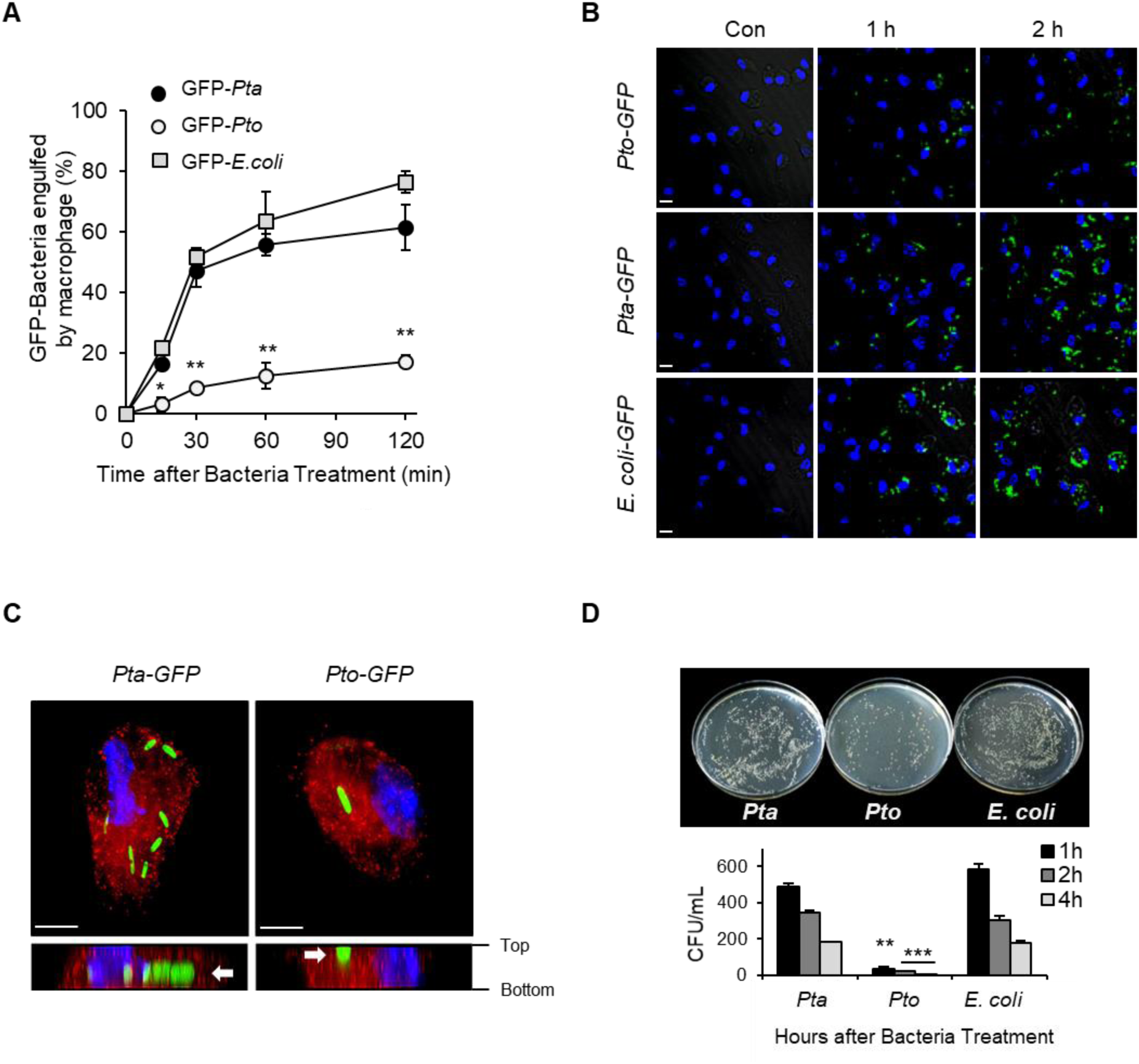
A plant pathogen, *Pseudomonas syringae* pv. Tomato, avoids phagocytic engulfment by mouse peritoneal macrophages. (*A*) Peritoneal macrophages from C57BL/6 mice were infected with GFP-expressing *Pta*, *Pto*, or *E. coli* [multiplicity of infection (MOI), 20:1] for the indicated times. The macrophages were analyzed by flow cytometry. Data are expressed as the mean ± SD, calculated from three different wells per time-point (\**P* < 0.05, \*\**P* < 0.01, compared with *Pta*). (*B*) Phagocytosis by mouse peritoneal macrophages was determined by immunofluorescence analysis. GFP-expressing *Pto, Pta,* or *E. coli* were fixed and analyzed by confocal laser microscopy (LSM510; Carl Zeiss). Scale bars, 20 μm. (*C*) Peritoneal macrophages were infected with GFP-expressing *Pto* or *Pta* (MOI, 20:1) for 30 min and then washed and fixed. Cells were then stained with a primary antibody against Rab5, followed by an Alexa Fluor 555-conjugated secondary antibody. All images were analyzed by confocal microscopy. (*D*) Internalized bacteria were detected in a gentamicin protection assay. Bacterial replication was expressed in terms of colony forming units (CFUs). Data are expressed as the mean ± s.d., calculated from three different plates per time point (\*\**P* < 0.01, \*\*\**P* < 0.001, compared with *Pta*).

Since phagocytosis involves not only internalization of bacteria but also phagosome maturation (*7, 8*), additional assays to confirm phagosome maturation were performed by measuring elimination of internalized bacteria and expression of phagosomal markers in macrophages. Although the numbers of internalized *Pto* bacteria were lower than the numbers of internalized *Pta* and *E. coli*, all internalized bacteria belonging to each of the three species were eliminated after incubation within macrophages, indicating cell lysis in the phagosome (Fig. S1*A*). No differences were observed in expression of early (Eea1 and Rab5) or late (Rab7 and Lamp1) phagosomal markers, indicating equivalent phagosome maturation in response to infection with each of the three species (Fig. S1*B*) (*27, 28*). These results indicate that during infection by *Pto*, phagocytosis by macrophages was inhibited at an early step (internalization) rather than at a late step (phagosome maturation).

### Bacterial type III secretion plays a pivotal role in inhibiting phagocytosis both *in vitro* and *in vivo*

Many pathogenic Gram-negative bacteria, including *Pto*, use type III secretion systems (T3SS) to secrete effector proteins directly from the bacterial cell into the host cytosol. In most cases, these effectors disturb or inhibit host defense mechanisms (*29*). Since the plant pathogen *Pto* inhibited the early stage of phagocytosis by macrophages, we hypothesized that T3SS-derived effectors play a pivotal role in suppression of phagocytosis. To verify this hypothesis, *Pto* Δ*hrpL*, a absence of regulator of effector genes (T3SS-null mutant) (*30*), was employed. As expected, *Pto* Δ*hrpL* showed recovery of phagocytosis determined by flow cytometry (Fig. 2*A*), gentamycin protection assay (Fig. 2*B*), and confocal microscopy (Fig. 2*C*). This observation was reproduced in a macrophage cell line, RAW 264.7 cells (Fig. S2*A*). The significant changes in expression of phagosome markers were not observed after infection with *hrpL*-deleted but not wild-type (WT) *Pto* (Fig. S2*B*). Hence, secretion of effectors via the T3SS controls phagocytosis at an early step: internalization of bacteria.

**Fig. 2.**
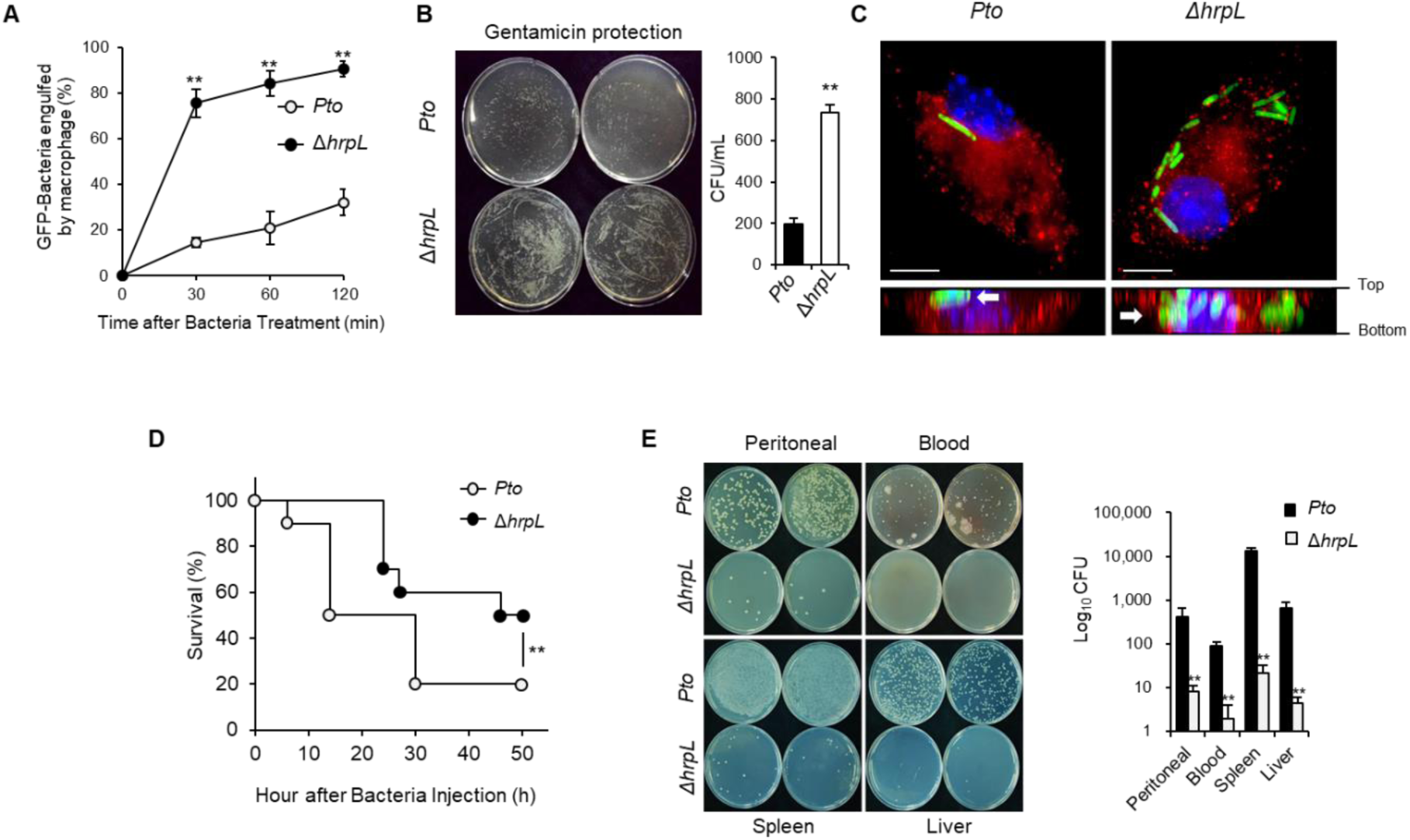
Deletion of the type III secretion system restored engulfment of mouse macrophages. (*A*) Peritoneal macrophages were infected with GFP-expressing *Pto* or Δ*hrpL* (MOI, 20:1) for the indicated periods, washed, scraped from the plate, and then fixed. Retained bacteria were analyzed by flow cytometry. Data are expressed as the mean ± SD, calculated from three different wells per time point (\*\**P* < 0.01). (*B*) Mouse peritoneal macrophages were infected with *Pto* or Δ*hrpL*, and bacterial replication was measured in a gentamicin protection assay. Data are expressed as the mean ± SD, calculated from three different plates per time point (\*\**P* < 0.01). (*C*) Peritoneal macrophages were infected with GFP-expressing *Pto* or Δ*hrpL* (MOI, 20:1) for 30 min and then washed, fixed, and stained with a primary antibody against Rab5, followed by an Alexa Fluor 555-conjugated secondary antibody. All images were generated by confocal microscopy. (*D*) *Pto* or Δ*hrpL* (108 CFU of live bacteria) was injected i.p. into C57BL/6 mice (n = 10). Viability was then assessed every 2 h (**P < 0.01, compared with *Pto*). (*E*) *Pto* or Δ*hrpL* was injected i.p. into C57BL/6 mice (n = 5). After 24 h, samples from the peritoneal cavity, blood, spleen, and liver were plated on agar and incubated for 24 h. Bacterial replication was measured by counting the number of CFUs. Data are expressed as the mean ± SD, calculated from three different plates per time point (**P < 0.01, compared with *Pto*).

Since the plant pathogen *Pto* was able to regulate phagocytosis in murine macrophages, the susceptibility of mice to *Pto* infection was investigated to determine whether a plant pathogen could cause sepsis in a different host kingdom. Intraperitoneal injection of *Pto* WT and Δ*hrpL* caused death within 50 h of observation. Interestingly, mice injected with *Pto* Δ*hrpL* showed a higher survival rate, with about 50% surviving at 50 h, while injection of *Pto* WT resulted in a 20% survival rate at the same time point (Fig. 2*D*). At 24 h after injection, the numbers of bacterial cells in the peritoneal cavity, blood, spleen, and liver were markedly lower in mice challenged with *Pto* Δ*hrpL* than in WT mice (Fig. 2*E*). Likewise, terminal deoxynucleotidyl transferase dUTP nick-end labeling (TUNEL) assays and hematoxylin and eosin (H&E) staining showed that the number of apoptotic cells in the liver, spleen, and lung following infection with *Pto* Δ*hrpL* was significantly lower than that following infection with *Pto* WT (Fig. S3*A*). Aspartate transaminase (AST) and alanine transaminase (ALT) levels in blood samples revealed a lower level of hepatic damage in *Pto* Δ*hrpL*-infected mice (Fig. S3*B*). Taken together, these results suggest that effectors secreted via the T3SS successfully inhibit phagocytosis *in vivo*.

### Type III effectors inhibit actin-mediated early phagosome formation

In the animal immune system, macrophages act as gatekeepers to prevent bacterial infection; however, bacteria have evolved various mechanisms to escape capturing by these cells (*16, 31*). Some animal pathogens produce actin-related proteins to regulate actin polymerization (*32, 33*), which is important for phagocytosis (*34, 35*). We showed that T3SS-derived effectors inhibit internalization of *Pto* via suppressing phagocytosis by macrophages; therefore, actin-related phagosome formation was further investigated. During *Pto* infection, F-actin accumulated in the cell membranes of peritoneal macrophages; however, after infection with *Pto* Δ*hrpL*, accumulation of F-actin was observed only around engulfed bacteria in the phagosome (total F-actin) (Fig. S4). To investigate this in greater detail, remodeling of RAW 264.7 macrophages through changes in the actin cytoskeleton during bacterial infection was examined. First, measurement of F-actin accumulation only in the phagosome (phagosomal F-actin) indicated that phagosome formation was restored in the absence of secretion of type III effector (Δ*hrpL*) (Fig. 3*A*). Second, time-lapse observation of RAW 264.7 macrophages during bacterial infection revealed that both the number and frequency of phagocytic cups were significantly higher in cells infected with Δ*hrpL* (Fig. 3*B* and Fig. S5). Additionally, compared with the Δ*hrpL* mutant, *Pto* markedly inhibited protrusion and retraction (Fig. 3C). Since the protrusion and retraction steps of cup formation are the critical steps in F-actin-mediated early phagosome formation (*9*), we examined phosphorylation of two key factors in actin regulation, LIM domain kinase 1 (LIMK1) and cofilin1. LIMK1 and cofilin1 are well-known regulators of transient actin polymerization and depolymerization (*36*). Interestingly, *Pto* infection induced marked phosphorylation of LIMK1 and cofilin1, while *Pto* Δ*hrpL* infection did not (Fig. 3*D*). LIMKs are serine/threonine kinases that are activated by other kinases through phosphorylation. Phosphorylated LIMK1 causes phosphorylation and subsequent inactivation of cofilin1, which regulates actin polymerization/depolymerization (*36*). These events inhibit phagocytic cup closure (*15*). Overall, these findings strongly suggest that inhibition of phagocytosis by T3SS-related effectors of *Pto* occurs through deregulation of actin polymerization/depolymerization induced by LIMK1/cofilin1 phosphorylation.

**Fig. 3.**
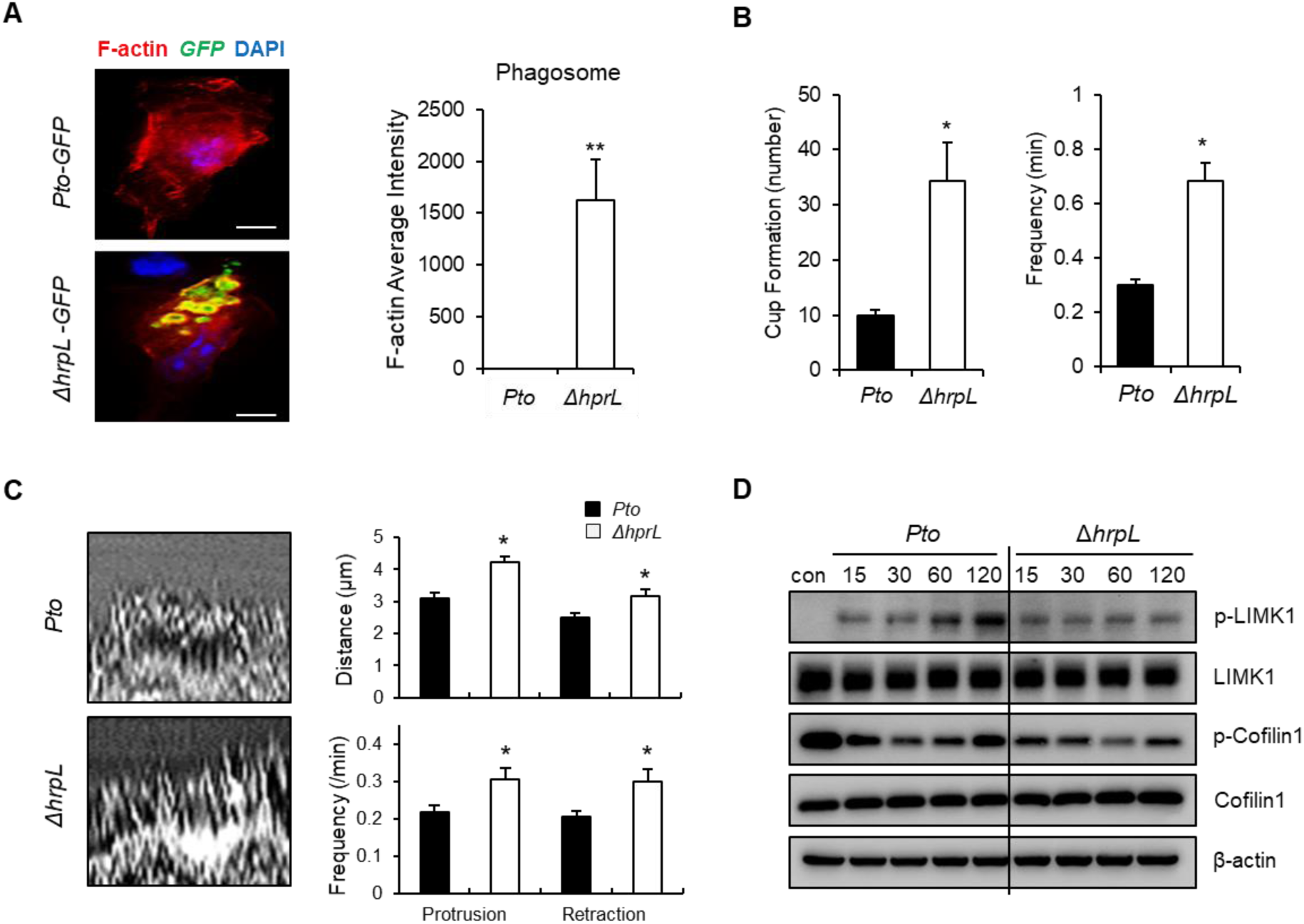
T3SS-mediated effectors of the plant pathogen *Pto* inhibit actin-mediated early phagosome formation. (*A*) RAW 264.7 cells were infected with GFP-labeled *Pto* or Δ*hrpL*. After 30 min, cells were fixed and stained with phalloidin to examine F-actin. The intensity of F-actin expression was measured using Metamorph software (n = 25). Scale bar, 5 μm (**P < 0.01, compared with *Pto*). (*B*) Temporal analysis of phagocytic cup formation by macrophages. Images were acquired at 15 sec intervals for 30 min after infection with *Pto*-GFP or Δ*hrpL*-GFP. For each group, phagocytic cup formation by at least 10 individual cells was monitored for 30 min, and the frequency was analyzed using Metamorph software (*P < 0.05, compared with *Pto*). (*C*) Deletion of *hprL* led to a notable increase in membrane ruffle formation. Lamella dynamics were analyzed using kymographs. For each group, membrane ruffles in at least 10 individual cells were monitored over a 15 min period by capturing digital images every 6 sec. Subsequently, three areas of interest were marked on each image using lines that crossed the cell lamella. Kymographs were then assembled using Metamorph software and ruffle formation was quantified. Data from five independent experiments are shown. (*D*) *P. fluorescens* (pHIR11) and HopQ1-expressing *P. fluorescens* (MOI, 20:1) were used to infect mouse macrophages at the indicated times and expression of LIMK1, p-LIMK1, p-cofilin1, cofilin1, and β-actin was examined by immunoblot analysis.

### The bacterial type III effector HopQ1 is central to inhibition of phagocytosis

*Pto* harbors at least 28 effectors that are fully active and expressed at levels detectable in a type III effector translocation assay in *Arabidopsis* and tomato plants (*37*). To identify the key effector protein(s), bacterial internalization via phagocytosis was evaluated in the *Pseudomonas fluorescens* EtHAn (Effector-to-Host Analyzer) strain (which is non-pathogenic but was engineered to express the structural T3SS) and in variants expressing 16 individual T3SS effectors. A gentamycin protection assay revealed that overexpression of HopQ1 clearly inhibits internalization of *P. fluorescens* EtHAn into macrophages (Fig. 4*A*). Additionally, *P. fluorescens* EtHAn expressing HopQ1 induced phosphorylation of LIMK1 and cofilin1 in macrophages, similarly to wild-type *Pto* infection (Fig. 4*B*). In addition, overexpression of HopQ1 in macrophages increased phosphorylation of LIMK1 and cofilin1 (Fig. 4*C*). To confirm translocation of HopQ1 from the bacterial cytosol to the animal cell, *Pto* WT, two T3SS-deficient mutants (Δ*hrcC* and Δ*hrpL*), and Δ28E (in which 28 effectors are knocked out) were transfected with a plasmid expressing a *hopQ1-cya* fusion protein and adenylate cyclase assays were performed. Cells infected with the two T3SS-deficient mutants showed little production of cAMP compared to cells infected with the other three bacterial strains (Δ28E, *Pto* WT, and *P. fluorescens* EtHAn), which indicates that translocation of HopQ1 was inhibited (Fig. S6, left two columns). By contrast, in the other three cell types in which type III secretion is intact, cAMP levels in macrophages were similar, suggesting successful translocation of HopQ1 from the bacteria to the animal cells (Fig. S6, right three columns). Therefore, HopQ1, which translocates into macrophages through the T3SS machinery, is critical for inhibition of phagocytosis.

**Fig. 4.**
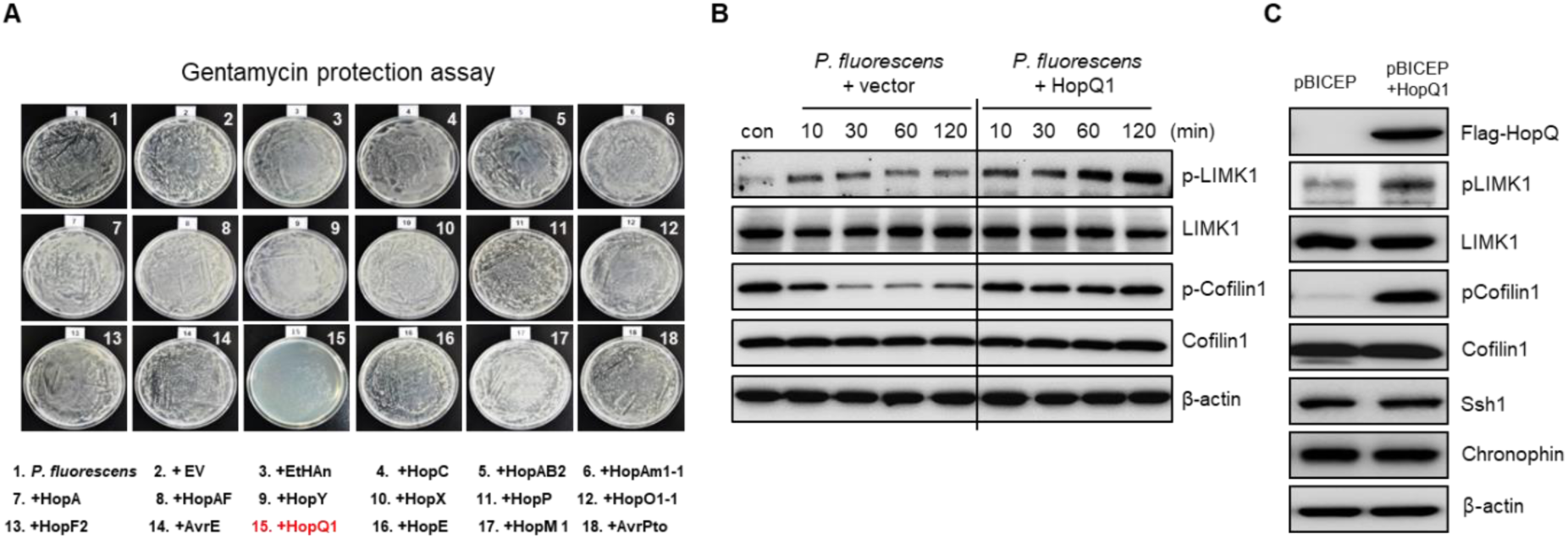
The T3SS-mediated effector HopQ1 activates LIMK1 and increases phosphorylation of cofilin *in vitro*. (*A*) Each effector protein of *P. syringae* tomato DC3000 was introduced separately into *P. fluorescens*. Macrophages were infected with each strain for 1 h at 37°C and bacterial replication was measured by determining the number of CFUs in a gentamicin protection assay. (*B*) Peritoneal macrophages were infected with GFP-expressing *Pto* or Δ*hrpL* (MOI, 20:1) for the indicated times and expression of LIMK1, p-LIMK1, p-cofilin1, cofilin1, and β-actin was examined by immunoblot analysis. (*C*) 293T cells were transfected with a control pBICEP vector or a pBICEP vector expressing HopQ1. At 24 h post-transfection, the levels of LIMK1, p-LIMK1, p-Cofilin1, Cofilin1, Ssh1, chronophin, and β-actin were determined by immunoblot analysis. Immunoblot analysis of HopQ1 expression was performed using an anti-Flag antibody.

### The effector HopQ1 physically interacts with the LIM2 domain of LIMK1

Various effector proteins secreted by *Pto* repress the plant defense response by targeting the proteins directly involved in signaling cascades related to gene regulation. To identify the target protein of HopQ1 that induces phosphorylation of LIMK1 and cofilin1, Flag-tagged HopQ1 was introduced into a kidney cell line (293T cells), and immunoprecipitation was performed. In this experiment, HopQ1 directly interacted with LIMK1, but not cofilin1 (Fig. 5*A*). Since LIMK1 is located upstream of cofilin1 in the actin-regulating cascade, these data demonstrate that deactivation of cofilin1 (phosphorylation) is due to phosphorylated LIMK1, which is induced by direct binding to the HopQ1 effector. Co-localization of HopQ1 and LIMK1 was confirmed via confocal microscopy (Fig. 5*B* and Fig. S7).

**Fig. 5.**
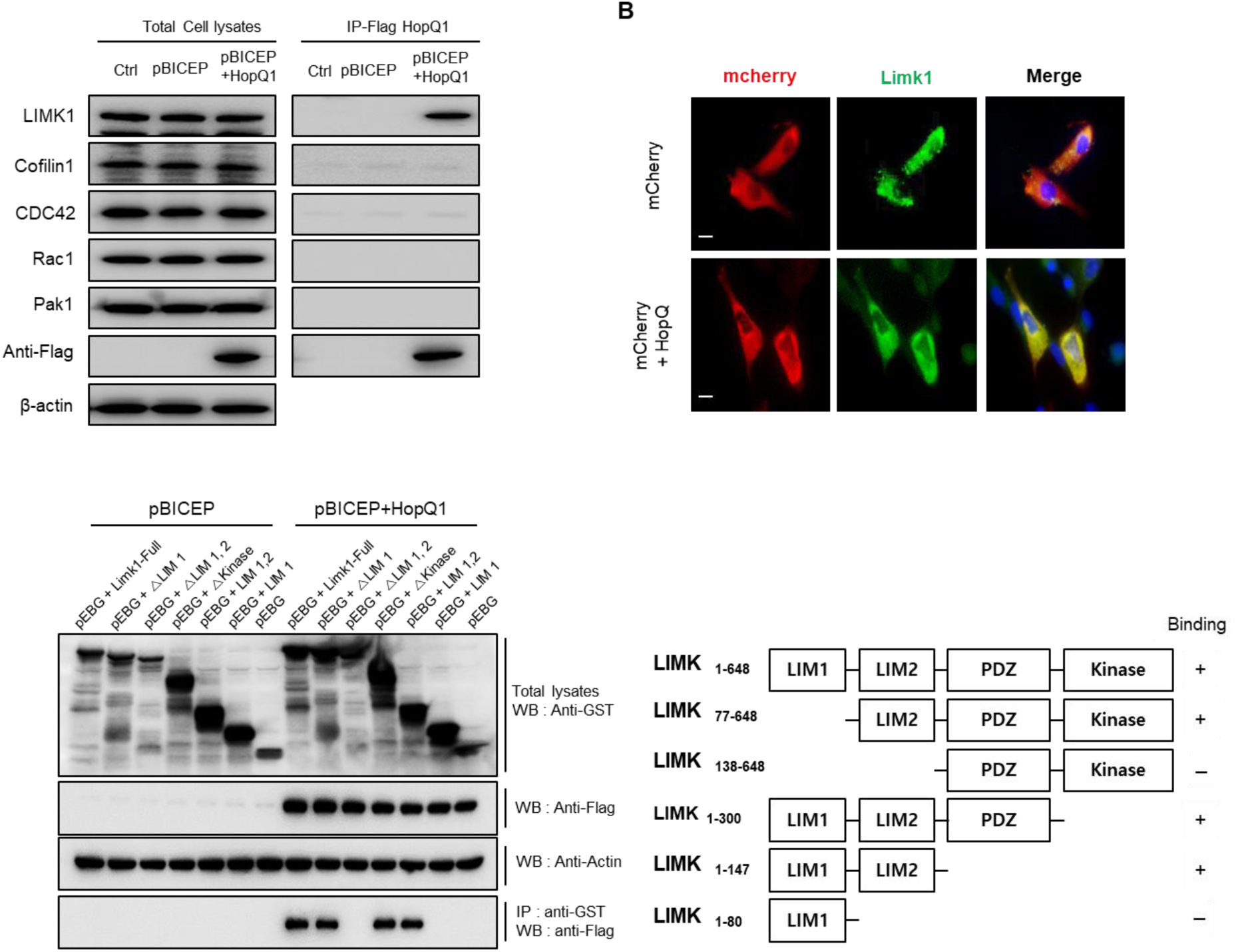
The effector HopQ1 interacts directly with LIMK1 in macrophages and inhibits phagocytosis. (*A*) 293T cells were transfected with a control pBICEP vector or a pBICEP vector expressing HopQ1. At 24 h post-transfection, cells were harvested and cell lysates were incubated for 2 h at 4°C with anti-DYKDDDDK (Flag)-tagged magnetic beads on a rotating apparatus. The interaction between HopQ1 and endogenous LIMK1, Cofilin1, Cdc42, Rac1, and Pak1 was examined by immunoprecipitation with an anti-Flag antibody, followed by immunoblot analysis. (*B*) 3T3-L1 cells were transfected with either GST-LIMK1 (green) or mCherry-HopQ1 (red) and then analyzed by confocal microscopy. Images show co-localization of LIMK1 and HopQ1 in 3T3-L1 cells. Scale bar, 10 μm. (*C*) HopQ1 interacted with LIMK1 via its N-terminal LIM2 domain. HopQ1 (Flag-expressing pBICEP)- and LIMK1 (GST-expressing pEBG)-encoding vectors were co-transfected into 293T cells and protein complexes were analyzed in a GST pull-down assay, followed by western blot analysis using anti-Flag and anti-GST antibodies, respectively.

LIMK1 contains a PDZ domain and a kinase domain (*38*). To examine how HopQ1 interacts with LIMK1, a series of LIMK1 deletion mutants lacking the LIM1, LIM2, PDZ, and/or kinase domains [ΔLIM1 (lacking residues 1–76), ΔLIM1, 2 (lacking residues 1–138), ΔKinase (lacking residues 301–648), ΔPDZ, Kinase (lacking residues 148–648), and ΔLIM2, PDZ, Kinase (lacking residues 81–648)] were constructed and their ability to interact with HopQ1 was tested in a glutathione S-transferase (GST) pull-down assay. Bands corresponding to HopQ1 were confirmed only in variants containing the LIM2 domain (specifically residues 84–138) (Fig 5*C*). Thus, HopQ1 interacts specifically with the LIM2 domain of LIMK1, possibly leading to inhibition of macrophage-mediated phagocytosis.

### Enhanced virulence of HopQ-transformed *Enterobacter cloacae* in mice

Next, *Enterobacter cloacae*, a T3SS-containing member of the normal gut flora of many mammals including humans (*39*), was used as a surrogate host to further examine the role of HopQ1 in phagocytosis. *E. cloacae* expressing *hopQ1* prevented phagocytosis by peritoneal macrophages (Fig 6*A*) and RAW 264.7 cells post-infection (Fig 6*B* and Movies S1-S2). Additionally, to confirm that HopQ1 regulates phagocytosis *in vivo*, mice were injected with *E. cloacae* expressing HopQ1 and the blood, spleen, and liver were examined after 24 h. Tissues from mice infected with HopQ1-expressing *E. cloacae* contained significantly more bacteria than those from mice infected with control *E. cloacae* (Fig 6*C*). Based on these findings, we propose a schematic model to describe the role of HpoQ1 in the immune response of macrophages (Fig 6*D*). In this model, HpoQ1 inhibits phagocytosis by interacting with LIMK1, the main regulator of cofilin. Phosphorylation of LIMK1 upon interaction with HopQ1 regulates the function of cofilin1, which is inactivated by phosphorylation. Thus, HpoQ1 ultimately prevents phagocytosis by disrupting F-actin accumulation in macrophages.

**Fig. 6.**
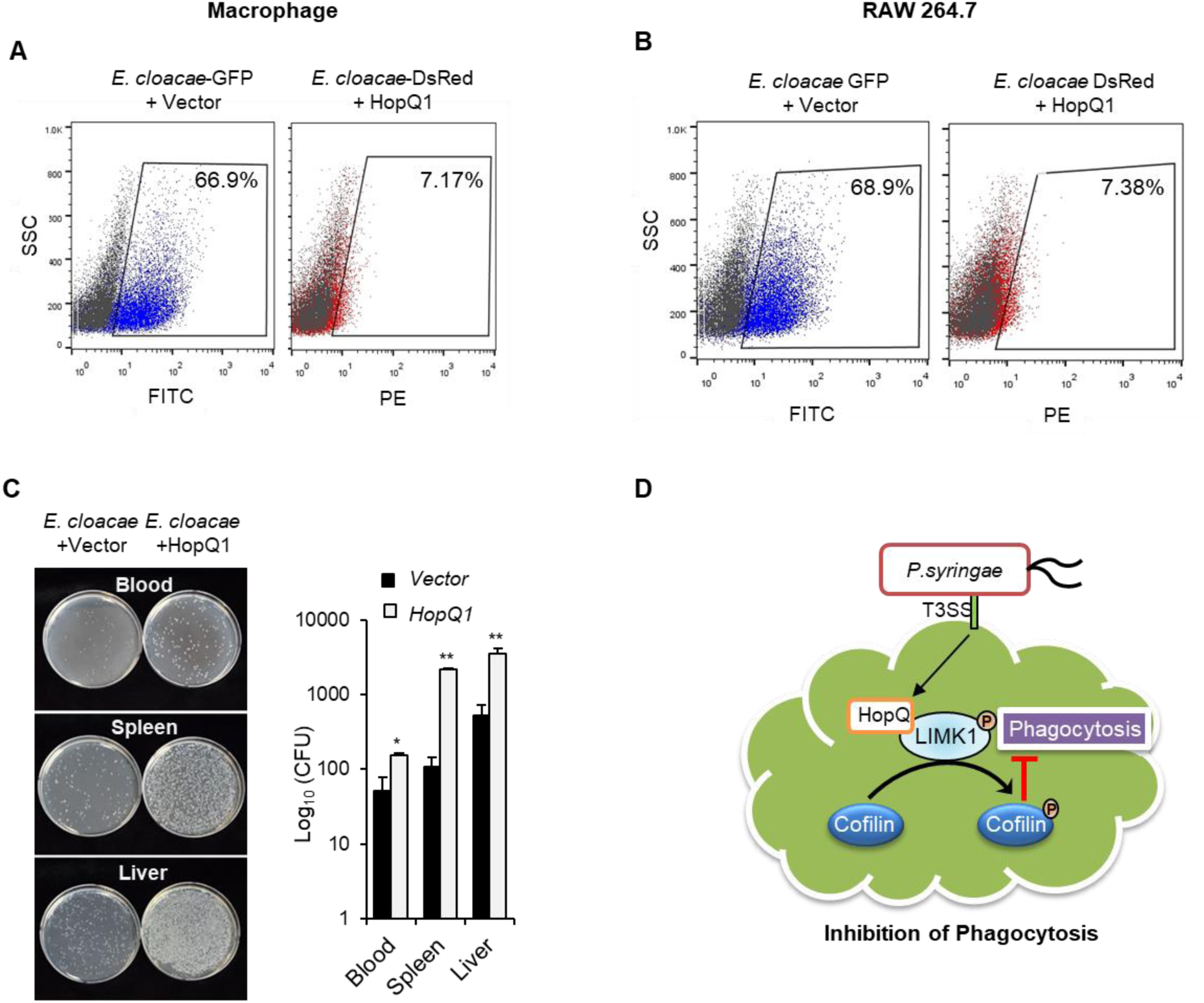
Phagocytic ability of macrophages infected with GFP-expressing *E. cloacae* or with DsRed- and HopQ1-expressing *E. cloacae*. (*A*) Mouse peritoneal macrophages were infected for 1 h with GFP-expressing *E. cloacae* or with DsRed- and HopQ-expressing *E. cloacae* and then rinsed. Bacterial elimination was examined by flow cytometry. (*B*) RAW 264.7 macrophages were infected with GFP-expressing *E. cloacae* or with both DsRed- and HopQ-expressing *E. cloacae* for 1 h and then rinsed. Flow cytometry analysis was used to confirm bacterial elimination. (*C*) *E. cloacae* (10^8^ CFU/mL) harboring a HopQ1 expression plasmid (or WT *E. cloacae*) was injected i.p. into C57BL/6 mice aged 10–12 weeks (n = 5). After 24 h, samples of blood, spleen, and liver were plated on agar and then incubated for 24 h. Bacterial replication was determined by counting the number of CFUs. Data are expressed as the mean ± SD, calculated from three different plates per time point (\**P* < 0.05, \*\**P* < 0.01, compared with *E. cloacae*). (*D*) A schematic model of the HopQ-mediated regulatory pathways involved in *P. syringae* pv. tomato-mediated inhibition of phagocytosis. *P. syringae* pv. tomato introduces effector proteins into host cells via the type III secretory system, thereby allowing the bacterium to avoid phagocytosis. This mechanism is mediated by the bacterial effector HopQ, which binds to the LIM domain of LIMK1 to activate the protein. Activated LIMK1 phosphorylates and thereby inhibits the function of cofilin1.

## DISCUSSION

Infection of animals by pathogenic plant bacteria is not rare. A previous study highlights the ability of specialized plant-associated bacteria to cause human and animal diseases and *vice versa* (*2*). Six genera (*Salmonella, Serratia, Enterobacter, Enterococcus, Pantoea,* and *Burkholderia*) caused symptoms in plants and mammals, either directly or after transmission by vectors such as insects (*2*). However, the mechanisms underlying evading animal innate immune system by plant pathogen have not been fully determined.

Phagocytosis by innate immune cells is a host defense response to infection, including microbial invasion. However, in some cases, phagocytosis can be impaired by effectors secreted by animal pathogenic bacteria. Possible mechanisms include direct disruption of the phagocytic machinery or resistance to the anti-microbial factors secreted by the phagocytes (*16, 27, 31*). For example, human pathogen *P. aeruginosa* disrupts the actin cytoskeleton by ExoT then inhibits internalization by macrophages (*40*). Outer membrane protein A of *E. coli* binds to complement regulatory proteins (C4b binding protein) and exhibits serum resistance (*41*). In addition, *S. aureus* presents complement inhibitor acting on C3 convertases and shows immune evasion (*42*).

Here, we provide evidence that plant pathogen-derived type III effectors can play a role in animal immune system. It has not been reported that plant pathogen *Pto* evades the animal innate immune system, phagocytosis. The mechanisms through which intracellular pathogens infiltrate host cells are well-studied (*16, 28*). In particular, regulation of actin dynamics is a well-known feature of the internalization of bacteria during phagocytosis (*9*), and some bacteria can escape from host cells or phagocytes by subverting host actin dynamics (*17, 31–33*). We demonstrated that formation of phagocytic cups increased markedly in macrophages infected with *Pto* lacking a T3SS (*Pto* Δ*hrpL*), suggesting that T3SS-related effectors of *Pto* may be involved in inhibiting the actin assembly network. Regulation of actin-filament network by *Pto* effector in a plant host is known as one of mechanism that disrupts plant immune system. One example is that HopG1 targets actin in the actin filament network of the plant cell (*43*).

The *Pto* effector HopQ1 played a key role in regulating LIMK1/cofilin1 phosphorylation, thereby inhibiting phagocytosis during infection. Originally, HopQ1 is the member of *hrp*-dependent outer protein (Hop) that injected into plant cells by the type III secretion system (*45*). The HopQ1 contains nucleoside hydrolase-like domain required for bacterial virulence (*46*). In plant host, phosphorylated HopQ1 interacts with 14-3-3 protein that controls functionally diverse signaling proteins (*47, 48*). Interestingly, in animal, 14-3-3 proteins contribute to the reorganization of the actin cytoskeleton by interacting with cofilin and its regulator LIMK149. These previous research might indicate interaction between plant HopQ1 and animal 14-3-3 proteins, thus phosphorylation of LIMK1 can be controlled by 14-3-3 interacting with HopQ1. However, our results support that physical interaction between HopQ1 and LIMK1. On the other hands, other Hrp outer proteins (e.g., HopF2, and HopAI1) interact with mitogen-activated protein kinase (MAPK) kinase (MAPKK), disrupting the MAPK cascade (*50, 51*). These indicate that Hop proteins family also can regulate kinase protein, directly. However, we did not demonstrate that how does HopQ1 phosphorylates the LIMK1 thus, further investigation is necessary.

In conclusion, the present study is the first to show that a plant pathogen can evade the mammalian innate immune system and can cause sepsis in animals. These studies also demonstrate that the effector of plant pathogenic bacterium, *Pto*, can translocate into macrophage cell and regulate the actin dynamics causing disruption of phagocytosis in different kingdom host. These results will facilitate further understanding of the molecular mechanism(s) underlying cross-kingdom pathogenicity. Causing sepsis by pathogenic bacteria presents different kingdom host rise the questions that how should we demonstrate the term ‘Infection’ and what is the Maginot line of host defense to determine in cross-kingdom pathogenicity? Pathogenic plant bacteria are potential reservoirs of human infection, which may have important implications for the emergence of infectious diseases. Moreover, continuous application of antibiotics results in drug-resistant pathogenic plant bacteria, such as pathovars of *P. syringae*, which may have a significant impact on human health.

## EXPERIMENTAL PROCEDURES

### Mice and bacterial infection

All animal-related procedures were reviewed and approved by the Institutional Animal Care and Use Committee (IACUC) of the Korea Research Institute of Bioscience and Biotechnology (KRIBB) and were performed in accordance with institutional (National Institutes of Health, USA) guidelines for animal care. C57BL/6 mice were purchased from Jackson Laboratories. The experimental groups were all age- and sex-matched. Mice were injected intraperitoneally (i.p.) with bacteria (1.0×10^8^ CFU) at 10–12 weeks of age. *P. syringae* pv. tomato DC3000 and *P. syringae* pv. tabaci were grown in King’s B medium for 24 h. To generate GFP-expressing bacteria, pathovars of *P. syringae* were transformed with pDSK-GFPuv (Km^R^).

### TUNEL assay, tissue staining, and blood analysis

Tissues were fixed in formalin and embedded in paraffin prior to sectioning (5 μm thick). Paraffin sections were then counterstained with H&E. The TUNEL assay was performed using the DeadEnd Colorimetric TUNEL System (Promega), according to the manufacturer’s instructions. Paraffin-embedded sections were deparaffinized in xylene for 5 min, rehydrated by immersion in a graded ethanol series, and fixed for 15 min in 4% paraformaldehyde in PBS. Tissue sections were permeabilized by incubation for 20 min in 100 μl of proteinase K solution (20 g/ml) at room temperature. All steps were performed according to the manufacturer’s instructions. Blood AST and ALT were analyzed using an automated blood chemistry analyzer (Hitachi 7150).

### Preparation of peritoneal macrophages and cell culture

Mouse peritoneal macrophages were harvested and cultured as described previously (*52, 53*). RAW 264.7 murine macrophages were purchased from the American Type Culture Collection and grown in RPMI 1640 medium (ThermoFisher Scientific/Gibco) containing 10% fetal bovine serum (FBS; HyClone). 293T cells were grown in DMEM (HyClone) supplemented with 10% FBS. 3T3-L1 cells were grown in DMEM supplemented with 10% fetal calf serum and antibiotics. The absence/presence of Mycoplasma was determined by PCR. All cell lines were free of Mycoplasma.

### Bacterial strains, plasmids, and transformation

*P. syringae* pv. tomato (*Pto*) DC3000, the *P. syringae* pv. tomato (*Pto*) DC3000 Δ*hrpL* mutant, *P. syringae* pv. tabaci, and *P. fluorescens* were cultured in King’s B broth media at 28°C (S1 Table) (*54*). *E. coli* DH5α and *E. cloacae* were cultured in Lysogeny broth and nutrient broth, respectively, at 37°C. The pDSK-GFPuv vector (green fluorescence) was introduced into bacteria via electroporation (*55*). To generate electrocompetent *P. syringae* strains, overnight cultures of *P. syringae* (derived from a single colony) were diluted to an optical density at a wavelength of 600 nm (OD_600_) of 0.2 in King’s B broth medium (20 g proteose peptone No. 3, 10 mL glycerol, 1.5 g K2HPO4, and 1.5 g MgSO4-7H2O per L) and then grown to an OD_600_ of 0.4–0.6. The cells were harvested by centrifugation at 4°C, washed four times with equal volumes of 300 mM cold sucrose, and finally resuspended (at 1/500 of the original culture volume) in 300 mM cold sucrose. Plasmid was added to 1 µL (100 ng/μL) of electrocompetent cells in a 0.2 cm cuvette. Cells were subjected to electroporation at 2.5 kV, 25 F, and 200 Ω using a Gene Pulser (Bio-Rad Laboratories). Next, King’s B broth medium (1 mL) was added and the cells were incubated at 30°C for 2 h with shaking prior to plating on King’s B agar containing kanamycin (50 µg/mL). The *P. fluorescens* type III effector strains are described in Table S1. The pBBRBB-DsRED (red fluorescence) and pBICEP-hopQ1-1 vectors were introduced into *E. cloacae* by electroporation, replacing 300 mM sucrose with 10% glycerol.

### Cloning and transfection

The pEBG and pBICEP vectors (Sigma) were used to generate GST-LIMK1 and FLAG-HopQ1 in 293T cells. The following primers were used: mouse LIMK1 (full), forward 5’-GCGTGGATCCACTAGTATGAGGTTGACGCTACTTT-3’ (Spe I) and reverse 5’-TCTAGAGTCGCGGCCTCAGTCAGGGACCTCGGG-3’ (Not I); mouse LIMK1 (ΔLIM1), forward 5’-GCGTGGATCCACTAGTGGCGAGTCTTGCCACGG-3’ (Spe I) and reverse 5’-TCTAGAGTCGCGGCCTCAGTCAGGGACCTCGGG-3’ (Not I); mouse LIMK1 (ΔLIM2), forward 5’-GCGTGGATCCACTAGTGAACAGATCCTACCTGAC-3’ (Spe I) and reverse 5’-TCTAGAGTCGCGGCCTCAGTCAGGGACCTCGGG-3’ (Not I); mouse LIMK1 (ΔKinase), forward 5’-GCGTGGATCCACTAGTATGAGGTTGACGCTACTTT-3’ (Spe I) and reverse 5’-TCTAGAGTCGCGGCCGCCCAGCACTTCCCCATG-3’ (Not I); mouse LIMK1 (LIM1, 2), forward 5’-GCGTGGATCCACTAGTATGAGGTTGACGCTACTTT-3’ (Spe I) and reverse 5’-TCTAGAGTCGCGGCCGATGACTGGAGTTACCACG-3’ (Not I); mouse LIMK1 (LIM1), forward 5’-GCGTGGATCCACTAGTATGAGGTTGACGCTACTTT-3’ (Spe I) and reverse 5’-TCTAGAGTCGCGGCCGCAAGACTCGCCATAGCG-3’ (Not I); HopQ1 (full), forward 5’-GACAAGCTTGCGGCCCATCGTCCTATCACCGCA-3’ (Not I) and reverse 5’-ATGCCACCCGGGATCTCAATCTGGGGCTACCGT-3’ (BamHI). The pLVX-mCherry-N1 vector (Clontech) was used to generate mCherry-tagged HopQ1 in 3T3-L1 cells. The primer sequences were as follows: mCherry-HopQ1, forward 5’-GGACTCAGATCTCGAGATGCATCGTCCTATCACC-3’ (Xho I) and reverse 5’-CATGACCGGTGGATCATCTGGGGCTACCGTCGA-3’ (BamHI). All cloning was performed using the In-Fusion HD Cloning Kit (Clontech), as recommended by the manufacturer. Cells were plated in 6-well plates at a density of 1.0×10^6^ cells per well. After culture for 24 h at 37°C to adjust to the transfection conditions, cells were incubated for 24 h at 37°C with DNA-polymer complexes using the TransIT-X2 Dynamic Delivery System reagent (Mirus Bio).

### Flow cytometry analysis of phagocytosis

Peritoneal macrophages and RAW 264.7 cells were plated in 12-well plates at a density of 1.0×10^6^ cells per well and then infected with 2.0×10^7^ CFUs of each GFP-expressing bacterium. After the indicated times, the wells were washed three times with cold PBS to remove any remaining bacteria. Cells were then scraped from the wells and immediately analyzed in a FACSCanto II flow cytometer (BD Biosciences). In total, 10,000 events per sample were collected and data were analyzed using FlowJo software v10.1.

### Immunostaining

Cells (1.0×10^5^ cells per well) were plated on round glass cover slips in 24-well plates and infected with bacteria (2.0×10^6^ CFU). After infection, cells were washed with cold PBS and fixed for 15 min at room temperature in 4% paraformaldehyde in PBS. Nuclei were stained for 5 min at room temperature with 4’,6-diamidino-2-phenylindole (DAPI; ThermoFisher Scientific) and the cells were washed again with cold PBS. For staining with primary antibodies, cells were permeabilized for 10 min at room temperature in 0.2% Triton X-100 in PBS and then incubated at 4°C overnight with antibodies specific for Rab5 (Cell Signaling Technology) or GST (Santa Cruz Biotechnology). Cells were then washed with PBS and incubated with Alexa Fluor 488-conjugated donkey-anti-rabbit IgG (Abcam) for 2 h at room temperature. To stain F-actin, cells were incubated for 30 min at room temperature with Alexa Fluor 555-conjugated phalloidin (ThermoFisher Scientific). After bacterial infection, live cells were incubated for 30 min at room temperature with LysoTracker Red DND-99 (ThermoFisher Scientific). Images were obtained using an LSM510 confocal microscope (Carl Zeiss).

### Western blotting, immunoprecipitation, and GST pull-down assays

Western blot analyses of cell lysates were performed as described previously (*42, 56*). For western blot analysis, antibodies specific for p-JNK (4668), JNK (9258), pNF-κB p65 (3033), NF-κB p65 (4764), p-Erk1/2 (4370), Erk1/2 (9102), p-p38 (9215), p38 (9212), Eea1 (3288), Rab5 (3547), Rab7 (9367), and chronophin (4686) were purchased from Cell Signaling Technology. Antibodies against pleckstrin (sc-136042), Lamp1 (sc-19992), and β-actin (sc-47778) were purchased from Santa Cruz Biotechnology. The anti-Flag antibody was purchased from Sigma. For immunoprecipitation, cell lysates were incubated for 2 h at 4°C with anti-DYKDDDDK (Flag)-tagged magnetic beads (Clontech Laboratories) on a rotating apparatus. The beads were then washed three times with cold lysis buffer, followed by an equal amount of 2X Laemmli sample buffer. To identify binding partners, antibodies specific for Limk1 (3842), pLimk1 (3841), p-cofilin1 (3313), cofilin1 (5175), Ssh1 (13578), and Pak1 (2602) were purchased from Cell Signaling Technology. Antibodies specific for Cdc42 (sc-8401) and Rac1 (sc-217) were purchased from Santa Cruz Biotechnology. For the GST pull-down assay, cell lysates were incubated with glutathione MagBeads (GenScript) for 1 h at room temperature, followed by washing three times in cold lysis buffer. An equal amount of 2X Laemmli sample buffer was then added to the beads. Antibodies bound to HopQ1 were detected using an anti-Flag antibody (Sigma).

### ELISA

Cytokines (TNF-α, IL-1β, and IL-6) in the culture medium were measured using DuoSet antibody pairs (R&D Systems). The color was developed with a 3,3’,5,5’-tetramethylbenzidine (TMB) substrate reagent set (BD Biosciences), and absorbance was measured at 450 nm using a microtiter plate reader (Emax; Molecular Devices).

### Gentamicin protection assay

Macrophages were inoculated with each bacterium for 1 h, treated with gentamicin for 1 h, and then washed with cold PBS to remove adherent bacteria from the cell surface. The cells were lysed in PBS containing 1.0% Triton X-100 for 15 min at room temperature. The lysates were then plated on agar plates and incubated for 20 h at 37°C, or in the case of plant bacteria at 30°C.

### Phagocytic cup formation

RAW 264.7 cells were seeded in Ibidi high 35 mm μ-dishes (Ibidi) at a density of 1×10^5^ cells/dish. After 24 h, cells were incubated with fresh culture medium and transferred to a live cell incubating chamber (Live Cell Instrument) under an atmosphere of 5% CO_2_ at 37°C. The chamber was set on the stage of an inverted fluorescence microscope (IX81-ZDC; Olympus) fitted with a UPLSAPO 20X objective lens. Bacteria, either WT or Δ*hrpL*, were added to the culture dishes prior to live image analysis. Phagocytic cup formation in RAW 264.7 cells was monitored every 15 sec over a 30 min period. Data analysis was performed using MetaMorph software, version 7.1 (Universal Imaging).

### Statistical analysis

All data are expressed as the mean ± SD. Differences between averages were analyzed using a the Student’s t test. The significance of differences in survival was determined using a log-rank test.

## Author Contributions

S.-J.Y., Y.-J.P., and C.-M.R. designed, performed, and analyzed experiments. S.H.L., S.-H.L., and S.C. provided technical assistance with experiments. S.-H.L., and J.-K. M. carried out the animal studies. Y.-J.P., C.-M.R., and I.P.C. supervised the project. S.-J.Y., Y.-J.P., J.-S.K., and C.-M.R. wrote the manuscript.

## Acknowledgments

This work was supported by grants from the KRIBB Research Initiative Program, Republic of Korea, the Woo Jang-Choon Project (PJ002015) of the Rural Development Administration, and by the National Research Foundation (NRF), which is funded by the Korean government (MSIP) (NRF-2015M3A9E6028953).

## SUPPORTING INFORMATION

### SUPPORTING MOVIE LEGENDS

Movie S1. Mouse peritoneal macrophages were infected with GFP-expressing *E. cloacae*. Initiation of phagocytosis was observed approximately 5 min after cells were treated with bacteria.

Movie S2. Mouse peritoneal macrophages were infected with DsRed- and HopQ-expressing *E. cloacae*. Initiation of cell phagocytosis was observed approximately 5 min after the cells were treated with bacteria.

**Supporting Table S1.**
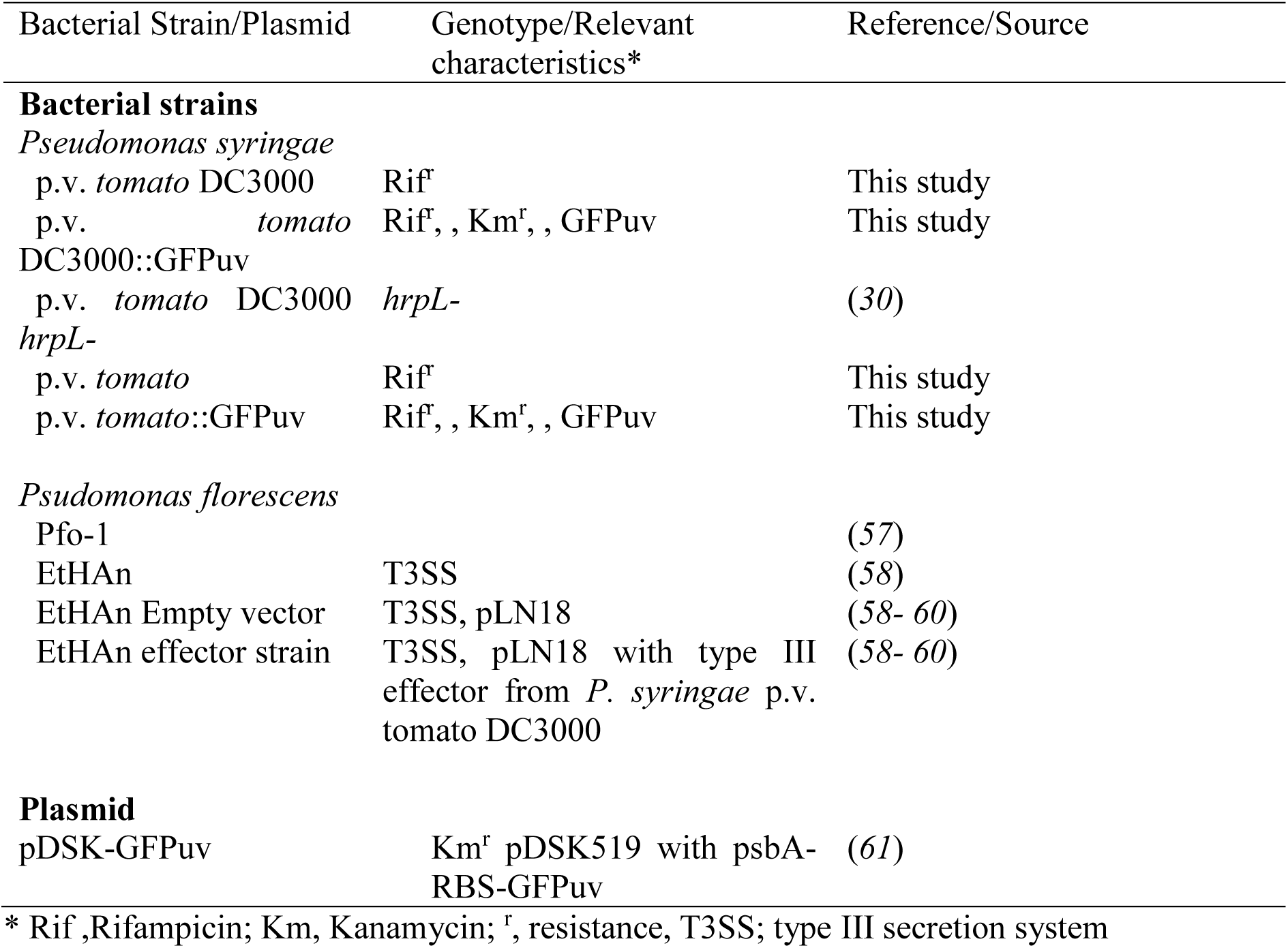
Bacterial stains and plasmids used in the study.

**Fig. S1.**
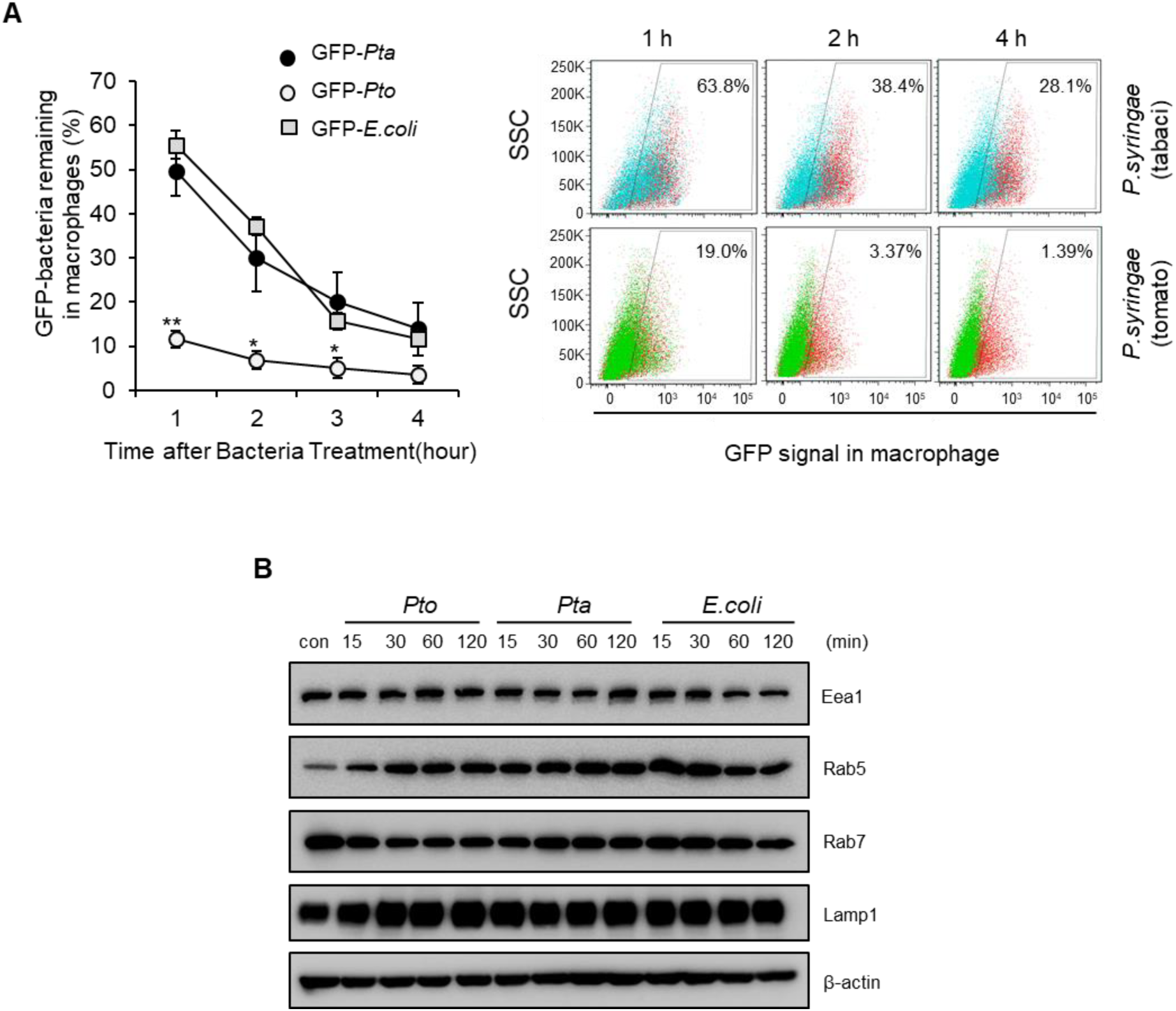
*P. syringae* tomato avoids engulfment during an early step in phagosome formation. (*A*) Mouse peritoneal macrophages were infected with *Pto-*GFP*, Pta*-GFP, or *E. coli*-GFP for 1 h and the remaining intracellular bacteria were detected by flow cytometry at the indicated times. Data are expressed as the mean ± s.d., calculated from three wells per time point (*P < 0.05, **P < 0.01, compared with *Pta*; left panel). Representative flow cytometry data is shown in the right panel. (*B*) Peritoneal macrophages were treated with *Pto*, *Pta*, or *E. coli* at an MOI of 20 and then harvested at the indicated times. Immunoblot analysis was performed to examine expression of Eea1, Rab5, Rab7, Lamp1, and β-actin.

**Fig. S2.**
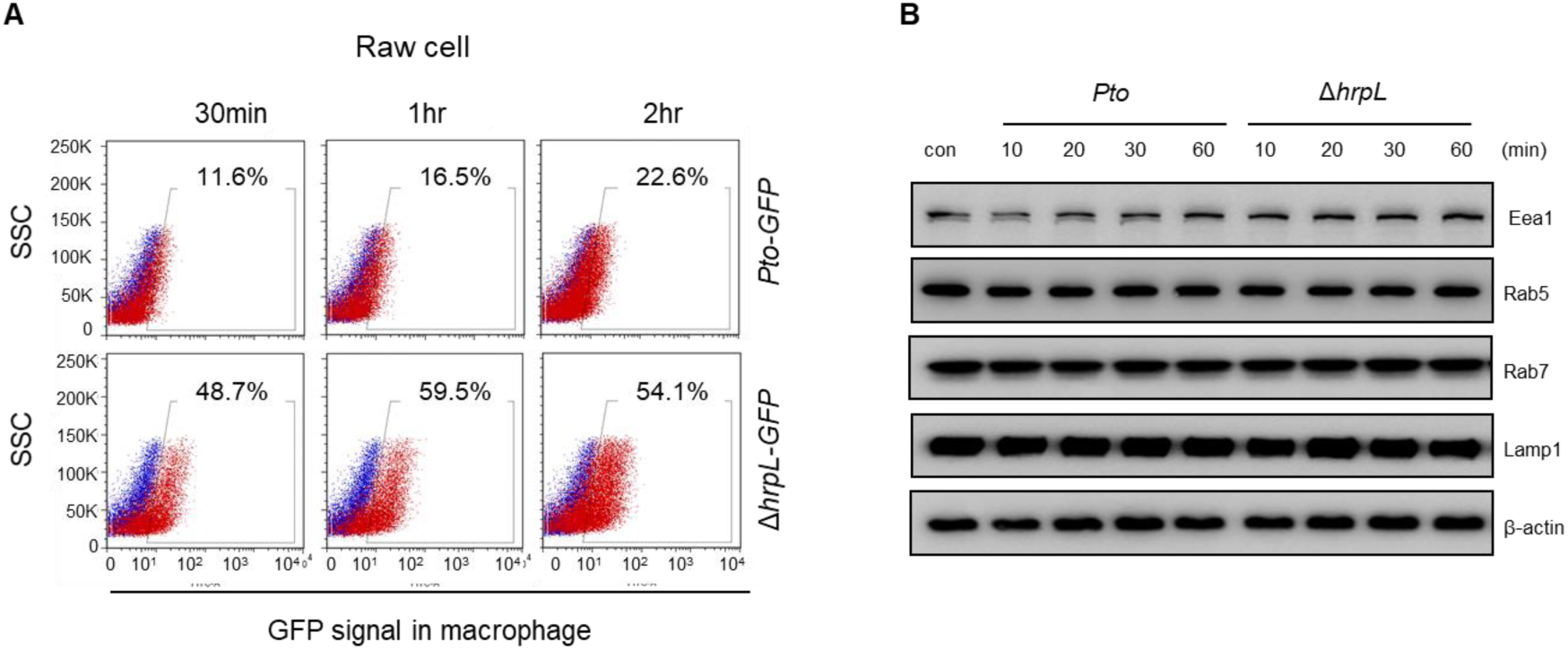
The *Pto* Δ*hrpL* mutant lacking the type III secretion system was efficiently engulfed by RAW 264.7 cells. (*A*) RAW 264.7 cells were infected with GFP-expressing *Pto* DC3000 or Δ*hrpL* (MOI, 20:1) for the indicated times and then washed, scraped from the plate, and fixed. Intracellular bacteria were analyzed by flow cytometry. (*B*) Peritoneal macrophages were infected with either *Pto* DC3000 or Δ*hrpL* (MOI of 20) at the indicated times and then analyzed by immunoblotting with antibodies against markers of early phagomes (Eea1, pleckstrin and Rab5), markers of late phagosomes (Rab7 and Lamp1), and β-actin (loading control).

**Fig. S3.**
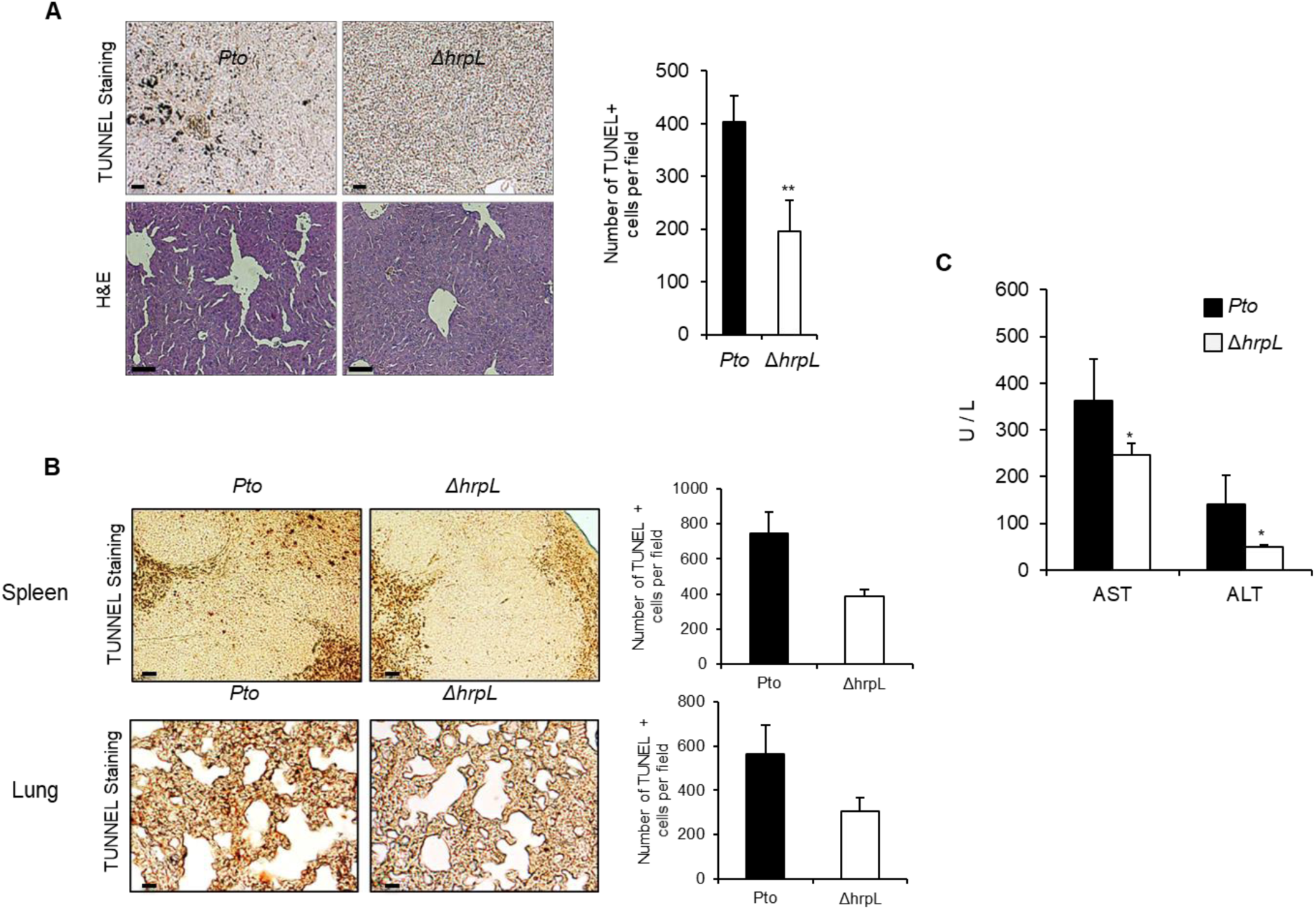
The *Pto* type III secretion system induces tissue damage *in vivo*. (*A*) *Pto* or Δ*hrpL* was injected i.p. into C57BL/6 mice aged 10–12 weeks (n = 5). Tissue samples from mice at 24 h post-bacterial injection were subjected to TUNEL assays and H&E staining. (*B*) TUNEL assays were performed on sections of spleen and lung obtained at 24 h post-bacterial injection. Apoptotic cells were counted in five randomly selected fields. Scale bar, 100 μm. Quantitation of the TUNEL staining is shown (\*\**P* < 0.01, compared with *Pto*). Scale bar, 100 μm. (*C*) At 24 h post-bacterial infection, AST and ALT levels in the serum were measured (\**P* < 0.05, \*\**P* < 0.01, compared with *Pto*).

**Fig. S4.**
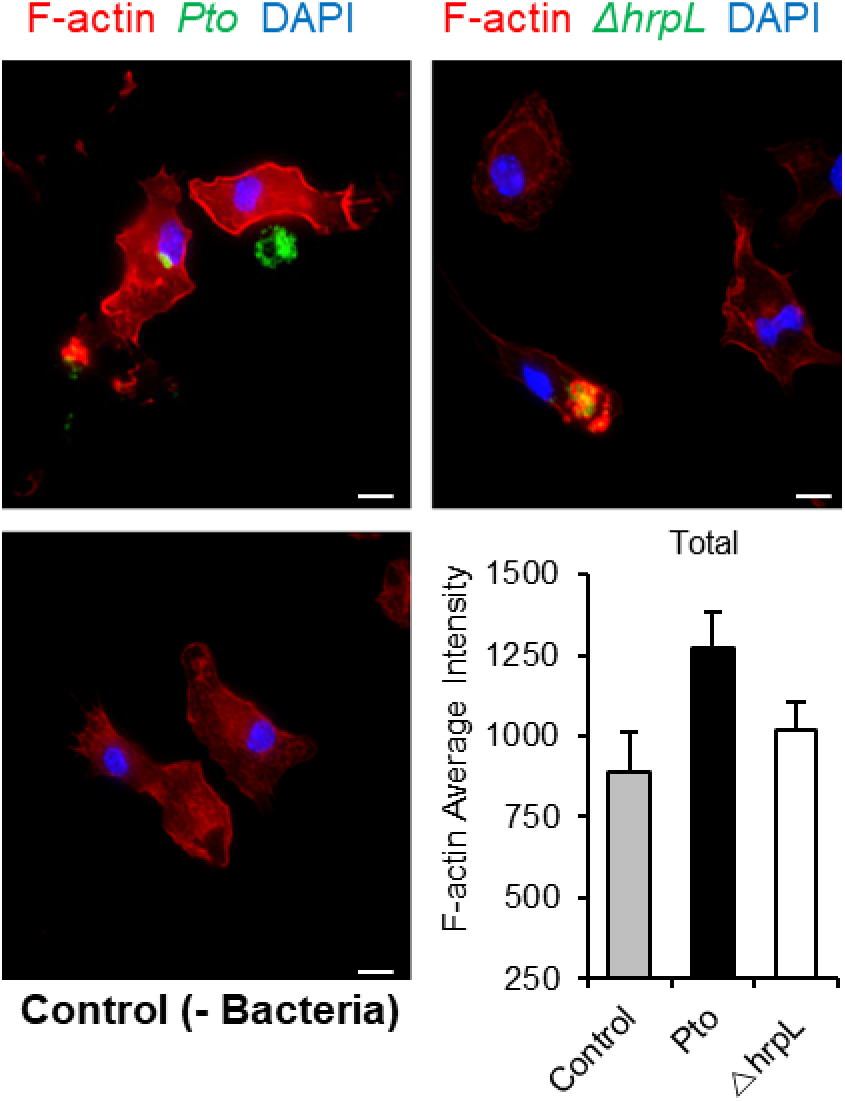
The intensity of F-actin formation. Peritoneal macrophages were infected with GFP-expressing *Pto* or Δ*hrpL* (MOI, 20:1) for 1 h and then washed, fixed, and permeabilized. Cells were then stained with Alexa Fluor 555-conjugated phalloidin for 30 min to examine F-actin formation. The intensity of F-actin formation was calculated according to cell size. Scale bar, 10 μm.

**Fig. S5.**
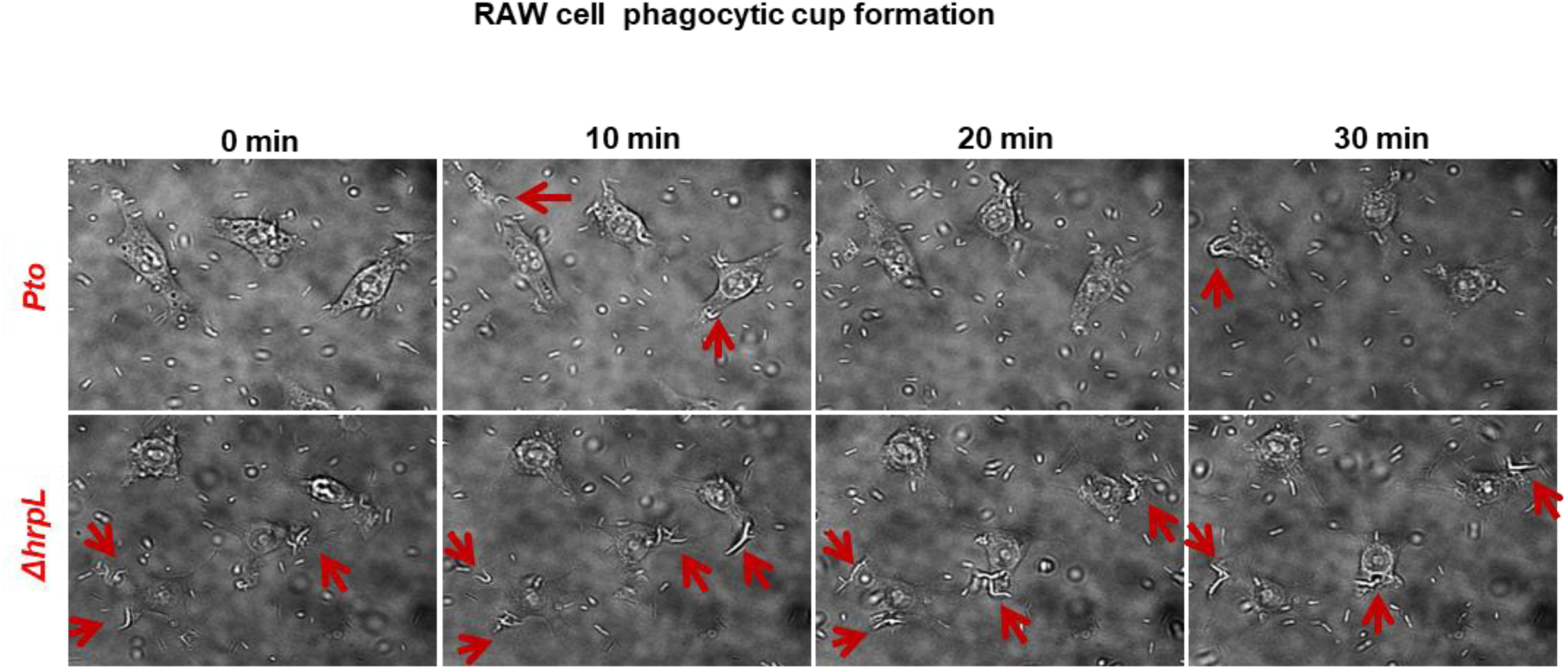
Phagocytosis of RAW 264.7 cells, as monitored by time-lapse microscopy. After treatment with each bacterium, images of live cells were captured every 15 sec for 30 min. Arrows indicate phagocytic cup formation. Scale bar, 20 μm.

**Fig. S6.**
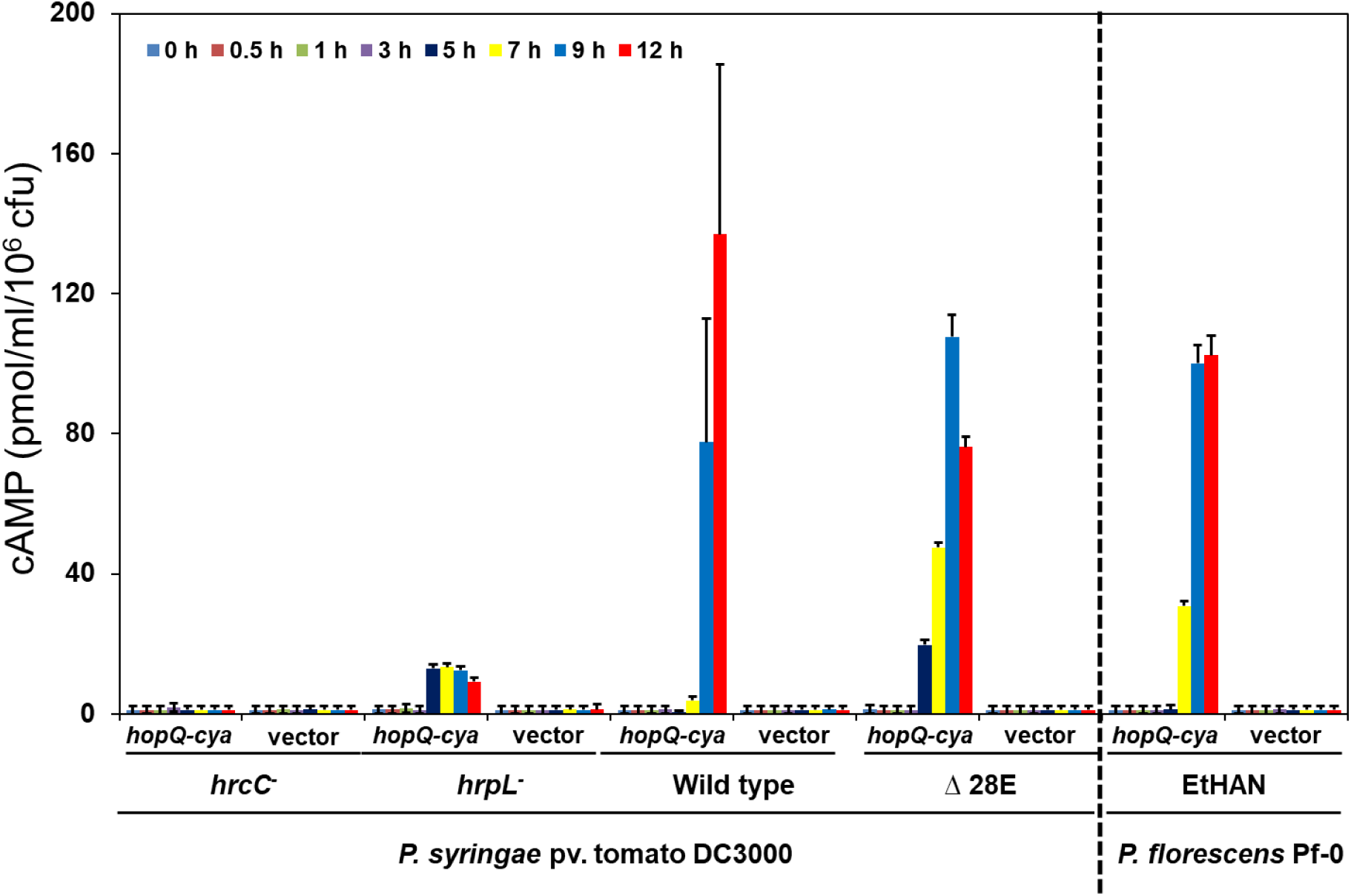
The effector HopQ1 translocates into macrophages through the T3SS machinery. RAW 264.7 macrophages were inoculated with *P. syringae* pv. tomato (wild-type and Δ*hrcC,* Δ*hrpL*, and Δ28E mutants) or *P. fluorescnes* EtHAn (Effector-to-Host Analyzer; a strain of *P. fluorescens* engineered to express an intact structural *Pto* T3SS) expressing *hopQ1*-*cya* or the control vector (pCPP3234). Accumulation of cAMP was measured at 0, 0.5, 1, 3, 5, 7, 9, and 14 h after bacterial inoculation. Results represent the mean ± SD. of triplicate wells for each sample. The graphs show data from one representative experiment. Repeat experiments on different days yielded similar results.

**Fig. S7.**
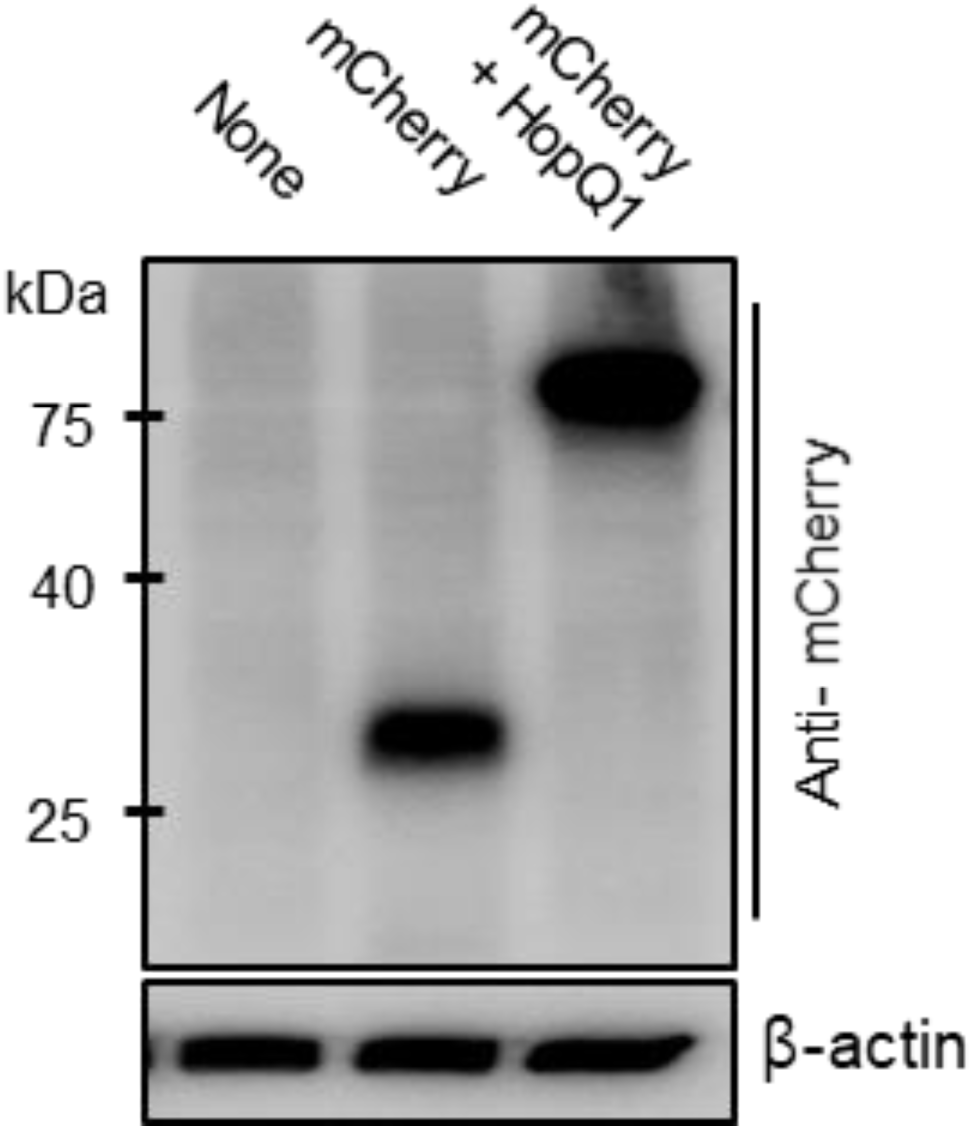
The level of mCherry-HopQ1 in 3T3L-1 cells. Expression of mCherry-HopQ1 was examined by immunoblot analysis.

